# Neutrophil-associated responses to *Vibrio cholerae* infection in a natural host model

**DOI:** 10.1101/2021.08.24.457598

**Authors:** Dustin Farr, Dhrubajyoti Nag, Walter J. Chazin, Simone Harrison, Ryan Thummel, Xixia Luo, Saumya Raychaudhuri, Jeffrey H. Withey

## Abstract

*Vibrio cholerae*, the cause of human cholera, is an aquatic bacterium found in association with a variety of animals in the environment, including many teleost fish species. *V. cholerae* infection induces a pro-inflammatory response followed by a non-inflammatory convalescent phase. Neutrophils are integral to this early immune response. However, the relationship between the neutrophil-associated protein calprotectin and *V. cholerae* has not been investigated, nor have the effects of limiting transition metals on *V. cholerae* growth. Zebrafish are useful as a natural *V. cholerae* model as the entire infectious cycle can be recapitulated in the presence of an intact intestinal microbiome and mature immune responses. Here, we demonstrate that zebrafish produce a significant neutrophil, IL-8, and calprotectin response following *V. cholerae* infection. Bacterial growth was completely inhibited by purified calprotectin protein or the chemical chelator TPEN, but growth was recovered by addition of transition metals zinc and manganese. Expression of downstream calprotectin targets also significantly increased in the zebrafish. These findings are the first to illuminate the role of calprotectin and nutritional immunity in combating *V. cholerae* infection. Inhibition of *V. cholerae* growth through metal limitation may provide new approaches in the development of anti-*V. cholerae* therapeutics. This study also establishes a major role for calprotectin in combating infectious diseases in zebrafish.

## Introduction

Cholera is a life-threatening severe diarrheal disease caused by the gram-negative bacterium *Vibrio cholerae. V. cholerae* is common in coastal regions and fresh, brackish or salt water, and cholera is endemic in many developing countries (1). These areas often have poor sanitation and infrastructure, aiding in the transmission of the bacteria via the ingestion of contaminated food or water through the fecal-oral route. While there are hundreds of *V. cholerae* serogroups in the aquatic environment, only O1 and O139 serogroups cause the pandemic cholera (2). The O1 serogroup can be further divided into classical and El Tor biotypes. While classical strains caused the first six pandemics, El Tor strains have caused the current pandemic, ongoing since 1961 (3).

Due to the extracellular nature of *V. cholerae* infection, cholera is generally considered to be a noninflammatory disease, though in reality some inflammation does occur. During early infection, acute cholera results in a pro-inflammatory response characterized by an increase in levels of inflammatory cytokines, receptors, and white blood cells (4, 5). Following this acute response is a noninflammatory convalescent phase, characterized by a suppression of inflammatory markers and an increase in vibriocidal IgM and mucosal antibody IgA (6-8). Innate immunity has been shown to be upregulated during *V. cholerae* infection, with neutrophils being documented as being an essential cell type. In a neonatal mouse model, neutrophils were shown to be recruited to site of infection and entering the lumen (9), while infection in neutropenic mice led to bacterial spread to extraintestinal organs and decreased survival, suggesting that the role of neutrophils in *V. cholerae* infection may be limiting the infection to the intestine (10). One major factor in neutrophil recruitment is the cytokine interleukin 8 (IL-8) (11). Known as neutrophil chemotactic factor, IL-8 functions to induce chemotaxis of neutrophils to the site of infection and stimulates phagocytosis. During cholera infection, intestinal epithelial cells have been shown to secrete IL-8 in response to *V. cholerae* flagellins as well as outer membrane protein OmpU (11, 12). In a zebrafish injury model, IL-8 was upregulated in response to acute inflammatory stimuli and was crucial for normal neutrophil recruitment to the site of injury and subsequent inflammation resolution (21).

One antimicrobial protein released by neutrophils is the heterodimer of EF-hand protein S100A8 and S100A9 termed calprotectin (CP). Released in neutrophil extracellular traps (NETs), as well as during cellular death, CP has been shown to make up 60% of the soluble protein in the cytosol of neutrophils (13). S100 proteins function as intracellular Ca^2+^ sensors, but when exported from the cytosol, CP functions to sequester transition metals including zinc, manganese, and copper (14, 51, 52). Thus, CP has antimicrobial activity by limiting access to these transition metals, a mechanism known as nutritional immunity (15). During *Candida albicans* infection, CP has been shown to be the major antifungal components of NETs (16) and can directly inhibit the growth several Gram negative and Gram positive bacteria (17). Increases in intestinal CP are not only seen in infectious diseases, as this protein is abundant in several inflammatory intestinal conditions, such as celiac disease (20), food allergies (19), and inflammatory bowel disease (IBD)(18). Many cases of these disorders can be diagnosed by detecting increases in fecal calprotectin levels. CP can also have downstream effects leading to the activation of a positive feedback loop of inflammation. Presence of CP can act as a cytokine and chemokine, can activate anti- and pro-inflammatory responses, and can also bind to receptors such as toll-like receptor 4 (TLR4) to activate a signaling cascade of inflammation through the transcription factor NF-κB (32, 33)

Zebrafish have previously been shown to be an effective model system for the study of *V. cholerae* infection, having many advantages over more commonly used mammalian animal models (3, 22-24). Among these advantages is that teleosts are a natural reservoir for *V. cholerae* species (25), the model is relatively inexpensive, easy to use, and recapitulates the characteristic diarrheal symptoms and transmission dynamics seen in humans (24). Colonization in the fish model also allows for the assessment of interaction between *V. cholerae* and the mature intact microbiota of the zebrafish (22). Furthermore, the zebrafish model remains relevant in studies examining the immune response due to the evolutionarily conserved nature of the immune system, allowing for direct parallels between the immune response of fish and humans (26).

While the zebrafish model has proven useful in the study of *V. cholerae*, no research has been published to date describing the zebrafish immune response to cholera infection, and our knowledge of the immune response of zebrafish and teleosts is still rudimentary and evolving (26). Moreover, our understanding of the immune response to *V. cholerae* continues to progress but remains far from complete. It will likely require innovative thinking and techniques to tease out the intricacies of this relationship. The link between the nutritional immunity protein CP and *V. cholerae* was not established until recently. To date, only one study examining adult cholera patients identified a 3.6-fold increase in S100A8 in lamina propria cells during acute-stage cholera; no other work has been published that further investigates this relationship (27).

In this study, using the zebrafish as a natural *V. cholerae* host model, we investigated the importance of neutrophils and the cytokine IL-8 during *V. cholerae* infection, and for the first time explore the relationship between the nutritional immunity protein CP and *V. cholerae*. Our results suggest that cholera induces a significant CP response, and that this protein and its ability to limit transition metals to inhibit bacterial growth may be an area of interest for the development of novel anti-*V. cholerae* therapeutics.

## Materials and Methods

### Bacterial strains and culture conditions

*V. cholerae* El Tor strain N16961 (Sm^r^ [100 µg/ml]), *V. cholerae* classical strain O395 (Sm^r^ [100 µg/ml]), *V. cholerae* environmental non-O1/O139 strain 25493 (Sm^r^ [100 µg/ml]), *V. cholerae* El Tor strain E7946 (Sm^r^ [100 µg/ml]), and *V. cholerae* non-O1/O139 strain AM-19226 (Sm^r^ [100 µg/ml]) were used in this study. Bacterial strains were frozen in 20% glycerol in Luria-Bertani (LB) broth (Difco, NJ, USA) at -80° C. For experimentation, each strain was then grown in LB broth (Difco, NJ, USA) at 37° C under shaking conditions (180 rpm) or on plates in LB agar (Difco, NJ, USA) with the appropriate antibiotic(s). Thiosulfate-citratebile-sucrose (TCBS) agar (Difco, NJ, USA) was used as selective media for *V. cholerae*.

### Zebrafish

Wild-type *AB* strain zebrafish were used for all experiments, except fluorescent microscopy experiments using Tg(*mpx:Dendra2*)^uwm4^/AB (53). For larval infections, zebrafish at 5 days post-fertilization (dpf) were used. Zebrafish were housed in an automated recirculating tank system (Aquaneering, CA, USA) using water filtered by reverse osmosis and maintained at pH 7.0 to 7.5. The tank water was conditioned with Instant Ocean salt (Aquarium Systems, OH, USA) to a conductivity of 600 to 700 S. Zebrafish were euthanized in 100 ml of 32-µg/ml Tricaine-S (tricaine methane sulfonate; MS-222 [Western Chemical, WA, USA]) for a minimum of 25-30 min after cessation of opercular movement.

### Adult zebrafish infection procedure

For experimental groups, 4-5 zebrafish were placed into a 400 ml beaker with perforated lids, containing 200 ml of tank water (autoclaved ddH2O with 60 mg/liter of Instant Ocean aquarium salts). Bacterial cultures were grown in LB broth at 37° C for 16 to 18 h with aeration. Bacteria was then washed once in phosphate-buffered saline (PBS) and diluted to a concentration of 10^9^ CFU/ml by measuring the OD at 600 nm. PBS diluted bacteria were then added directly to beakers to an infection concentration of 2.5 × 10^7^ and plated using serial dilutions for verification. Control fish were exposed to 1 ml of 1X PBS. Beakers containing fish were then placed in a glass-front incubator at 28° C for the duration of the experiment.

### Larval zebrafish infection procedure

5 day post-fertilization zebrafish larvae were placed into 50 ml beakers containing 20 ml sterilized tank water containing 2.5 × 10^7^ CFU/ml *V. cholerae*. For designated drug treatment experiments, the selective nonpeptide inhibitor SB 225002 (Tocris, Minneapolis, MN) was added at a concentration of 5 µM directly to beakers containing fish for 1 hr. prior to infection. After 24 hours, larvae were euthanized using tricaine solution, added directly to 300 ml trizol solution (Invitrogen, Waltham, MA) and homogenized using a pellet pestle (Fisher Scientific, Pittsburgh, PA). RNA was purified by ethanol precipitation and resuspended in RNase-free water. cDNA production and subsequent qRT-PCR were performed as described below.

### Intestinal colonization assessment

At specified time points, adult zebrafish were euthanized using tricaine. Intestines were aseptically removed and placed in homogenization tubes (2.0-ml screwcap tubes; Sarstedt, Nümbrecht, Germany) with 1.5 g of 1.0-mm glass beads (BioSpec Products, Inc., Bartlesville, OK) and 1 ml of 1 PBS, and held on ice. Homogenization tubes were loaded into a Mini-Beadbeater-24 (BioSpec Products, Inc.). Serial dilutions of homogenized tissue were plated onto LB agar plates with appropriate antibiotics.

### RNA isolation and qRT-PCR

Intestinal tissue from adult zebrafish was homogenized in 1ml 1X PBS using homogenization beads as described above. RNA was then extracted using Qiagen RNeasy Mini Kit (Qiagen, Hilden, Germany). Larval zebrafish were added directly to 300 ml trizol solution (Invitrogen, Waltham, MA) and homogenized using a pellet pestle (Fisher Scientific, Pittsburgh, PA). RNA was purified by ethanol precipitation and resuspended in RNase-free water. Total RNA was resuspended in RNase-free water and quantified using a NanoDrop. cDNA was then synthesized using Invitrogen SuperScript III First-Strand Synthesis System cDNA kit (Invitrogen, Waltham, MA). qRT-PCR was performed using SYBR green (Applied Biosystems, Foster City, CA). Quantification of gene expression was determined using the comparative ^ΔΔ^CT method. Gene expression was normalized to the endogenous reference β-actin level and was reported as fold change relative to the reference gene.

### Metal-reversible antimicrobial assays

To assess the metal binding and inhibitory properties of calprotectin and the zinc-chelator TPEN (Tocris, Minneapolis, MN) on *V. cholerae*, varying concentrations of the protein or chelator were added directly to LB broth media in a 96 well plate, incubated for one hour at 37° C, then bacteria was added and grown at 37° C with aeration. At the designated time points, the plate was then spectrophotometrically read at an O.D.600. Mn^2+^ and Zn^2+^ were then added to *V. cholerae* culture in increasing quantities to restore bacterial growth in calprotectin and TPEN experiments, and O.D.600 was read at 24 hrs. Recombinant wild-type and mutant CPs were expressed and purified as described previously (49).

### IL8 morpholino knockdown

Morpholino (MO) microinjections were performed as previous described (54). Briefly, one-to four-cell-stage wild-type (AB) embryos were microinjected with the morpholino solution at 4 ng/embryo. For this study two morpholinos were utilized, both obtained from Gene Tools (Philomath, OR): *cxcl8-ll* El/ll MO (Sequence 5’-GGTTTTGCATGTTCACTTACCTTCA-3’), which is a previously-established splice-blockingmorpholino for IL8 (21), and the Standard Control MO, which has no know target in zebrafish (55). At 48 hpf, embryos from each group were pooled separately and harvested for qRT-PCR, as described above.

### Imaging of infected zebrafish intestines

Fish were euthanized using tricaine after 24 hr infection, intestines were removed and then placed in 10% zinc formalin for 24 hrs. Next, intestines were placed in 70% ethanol and shipped to Reveal Biosciences (Reveal Biosciences, Inc., San Diego, CA) for fluorescent imaging as follows: samples were dewaxed in Xylene three times, then cleared using 100% alcohol twice. Samples were then hydrated in 95% alcohol and rinsed in distilled water twice. Slides were mounted using EMS, and stained using Fluoro-Gel II with DAPI and an anti-*V. cholerae* polyclonal antibody (KPL BacTrace) then counterstained with a secondary antibody conjugated to Alexa Fluor 647.

### Intestinal homogenate ELISA

Infection procedures were performed as described above. Intestinal tissue was removed and placed into 100 µL 1X PBS and homogenized using pellet pestles. Next, 50 µL of RIPA buffer was added and samples were centrifuged at 5000 × g for 5 minutes to remove debris. Samples were then diluted 1:25 with 1X PBS and fish calprotectin ELISA kit was ran according to manufacturer’s instructions (MyBioSource, San Diego, CA) as follows: 50 µL standard or sample were added to each well. 100 µL HRP-conjugate reagent was added to each well, and the plate was incubated for 60 minutes at 37° C. Wells were washed 4 times, 50 µL Chromogen solution A and B was added to each well and incubated for 15 minutes at 37° C. 50 µL stop solution was added to each well and after 15 minutes the plate was read spectrophotometrically at an O.D. 450 nm.

### Statistical analysis

Experiments were performed in triplicate on separate occasions, unless otherwise specified. Data shown are presented as the mean ± standard deviation (SD). All statistical analyses, t-tests and two-way ANOVAs were performed using Prism version 7.0 for Windows (GraphPad Software, La Jolla, CA).

### Ethics statement

All animal procedures were approved by the Wayne State University IACUC, protocol #18-10-0809. Zebrafish were euthanized in 100 ml of 32-µg/ml Tricaine-S (tricaine methane sulfonate; MS-222 [Western Chemical, WA, USA]) for a minimum of 25-30 min after cessation of opercular movement.

## Results

### In-vivo neutrophil response

Due to the extracellular nature of *V. cholerae* infection, we hypothesized that the innate immune system would respond early, with neutrophil recruitment into the lumen of the gut that peaked at roughly 24 h, as has been previously reported in the intestine (28). To assess the zebrafish immune response to *V. cholerae*, we inoculated fish via immersion using 2.5×10^7^ CFU of either pandemic O1 *V. cholerae* strain E7946 (labelled “El Tor”) or environmental *V. cholerae* strain 25493 (labelled “non-O1”). Fish were then incubated and sacrificed at 24 h post infection (h), 72 h, or 120 h. After euthanasia, RNA was isolated from fish intestines and qPCR was used to assess neutrophil (mpx) and IL-8 responses. In the El Tor infected fish, neutrophil gene expression was significantly increased at 24 h and 72 h, while it was significantly decreased at the 120 h time point (Fig. 1A). In the non-O1 strain infected fish, only 24 h fish had significantly increased levels of neutrophils as compared to control (Fig. 1B). The cytokine IL-8, which induces neutrophil chemotaxis and phagocytosis (21), was also significantly increased at all three time points in fish infected with either of the strains (Fig. 1C and 1D).

**Figure 1.**
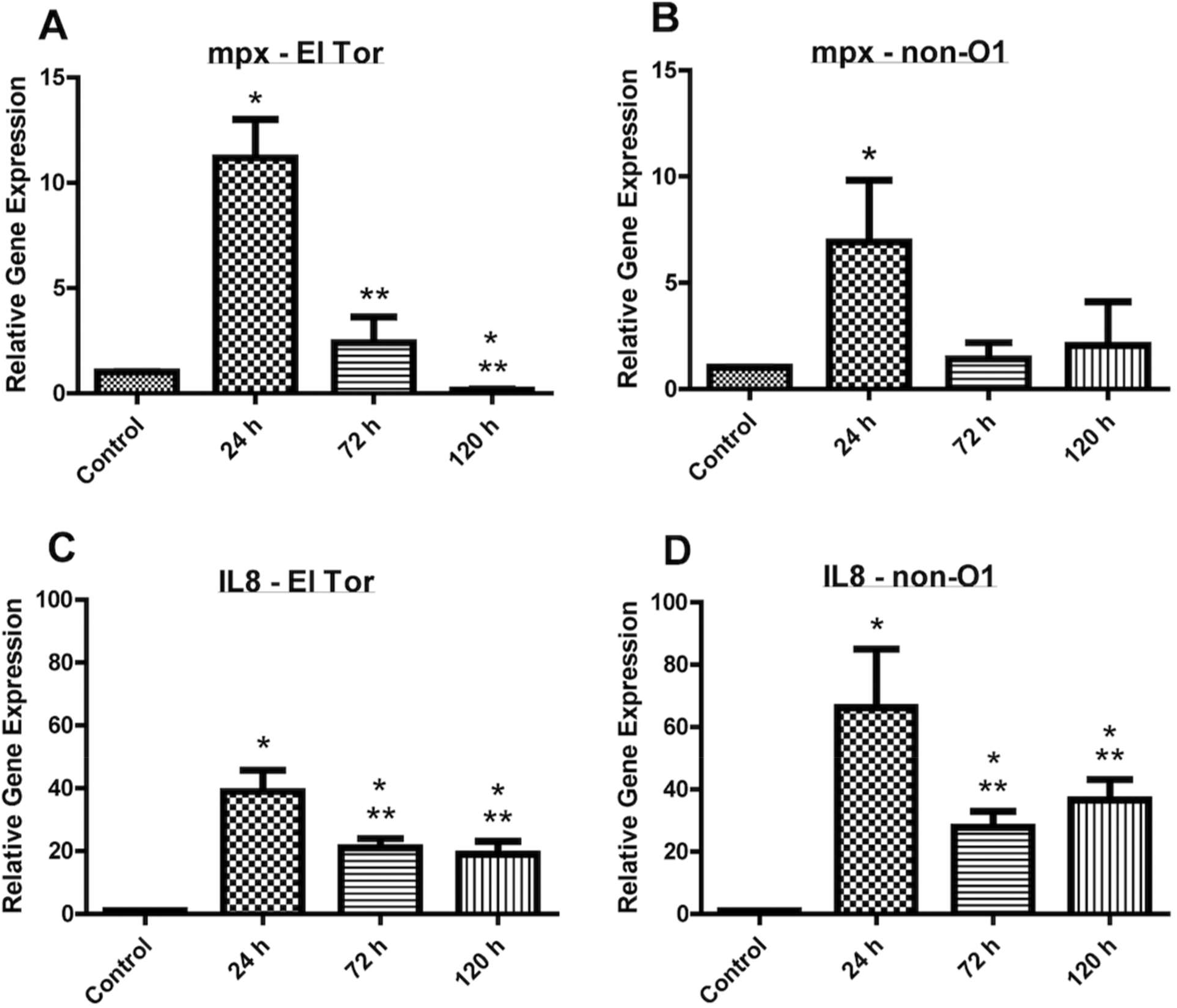
Neutrophil (mpx) and IL-8 (IL-8) levels increase during *V. cholerae* infection. WT zebrafish were infected with E7946 (El Tor) or 25493 (non-O1) strains of *V. cholerae* at 2.5 × 10^7^ CFU/mL and then sacrificed at the indicated time points. mRNA levels were determined through qRT-PCR. Gene expression was normalized against β-actin and expressed as fold change. Error bars indicate standard deviation. Data shown is from three independent experiments. *, P < 0.05 as compared to control, ** P < 0.05 as compared to 24 h infection.

Transgenic (mpx:dendra) zebrafish were also used to image neutrophil recruitment into the lumen of the gut at 24 h time points via fluorescent microscopy. A primary polyclonal antibody directed against *V. cholerae* along with a secondary monoclonal antibody with fluorescent tag (Alexa fluor 647) was used. These images show red fluorescent bacteria forming biofilms along villi projections of the intestinal epithelial cells (nuclei stained blue, DAPI) of the zebrafish, which has been previously reported as the site of *V. cholerae* biofilm formation in both mammals and zebrafish (24, 36). Within these areas of red fluorescence, individual green fluorescent neutrophils can be seen infiltrating bacterial biofilms, which can be seen in both El Tor and non-O1 infected fish (Fig. 2).

**Figure 2.**
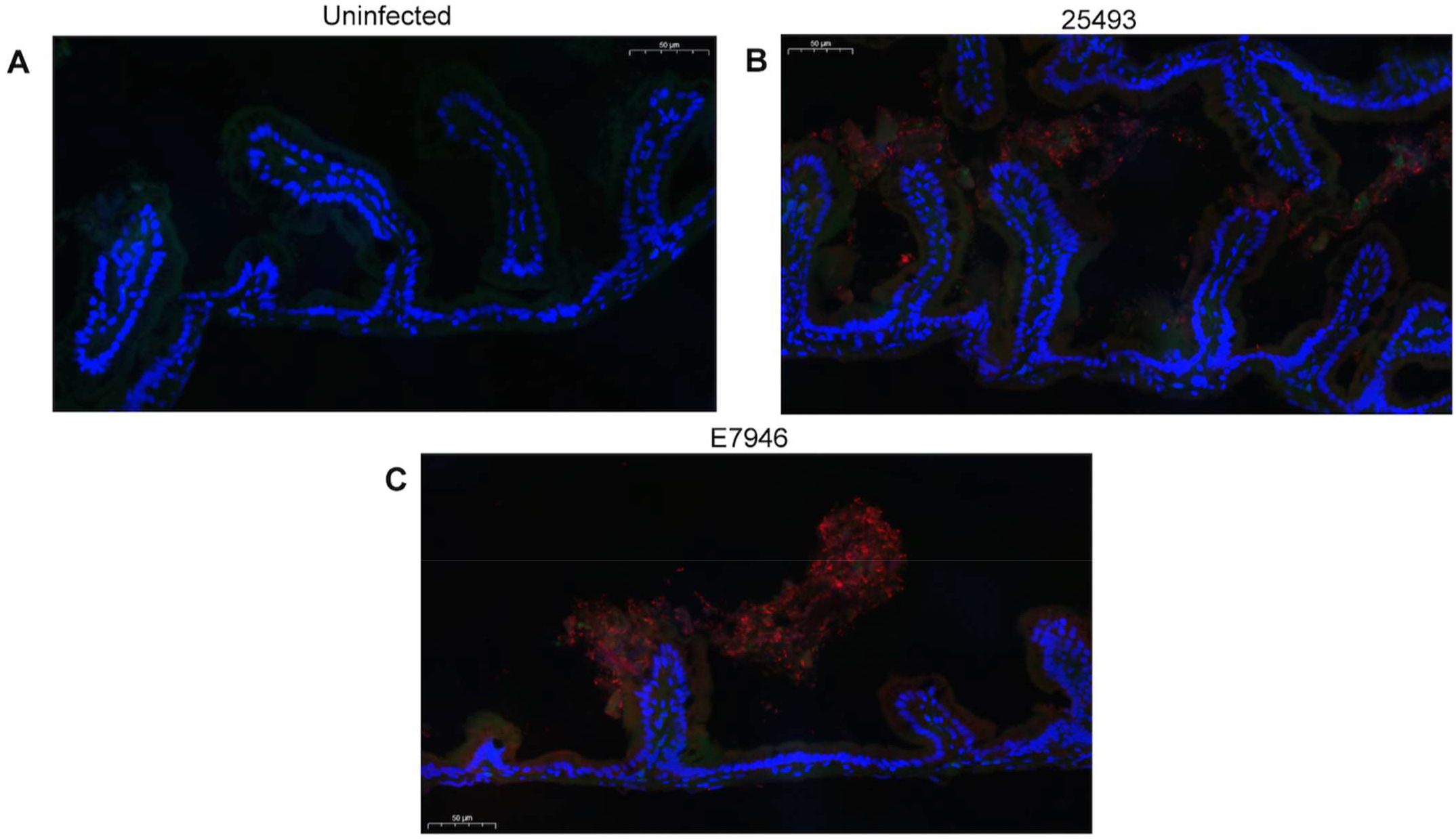
Fluorescent microscopy of neutrophil response to *V. cholerae* infection of transgenic (mpx:dendra) zebrafish intestinal epithelium. Fish were exposed to *V. cholerae* for 24 h then sacrificed, fixed, and prepared for sectioning. Bacteria were visualized using a polyclonal primary antibody against *V. cholerae* and a secondary antibody carrying a red fluorescent (Alexa fluor 647) tag. Blue fluorescence (DAPI) represents intestinal epithelial cell nuclei, red fluorescence (Alexa fluor 647) represents *V. cholerae* bacteria, and green fluorescence (dendra) represents neutrophils. (2A) Uninfected fish (2B) *V. cholerae* strain 25493 infected fish (2C*) V. cholerae* E7946 infected fish. Scale bar in all panels is 50 microns.

### In-vivo calprotectin response

We next wanted to investigate in-vivo levels of neutrophilic associated antimicrobial proteins in the fish intestine. RNA levels of the nutritional immunity protein calprotectin were measured at designated time points, with gene expression being significantly increased for both strains at 24 h and 72 h, but returning to basal levels by 120 h (Fig. 3A and 3B). Each timepoint was also significantly different from every other. We then wanted to determine if the in-vivo calprotectin response was seen during infection with other *V. cholerae* serogroups and strains. For these experiments, fish were infected with O395 classical, N16961 El Tor, AM-19226 non-O1/O139, and 25493 non-O1/O139 and euthanized at 24 h. In this experiment, only the environmental strain 25493 induced significantly increased levels of calprotectin RNA, therefore we decided to use this strain in later experiments that had reagent limitations (Fig. 3C).

**Figure 3.**
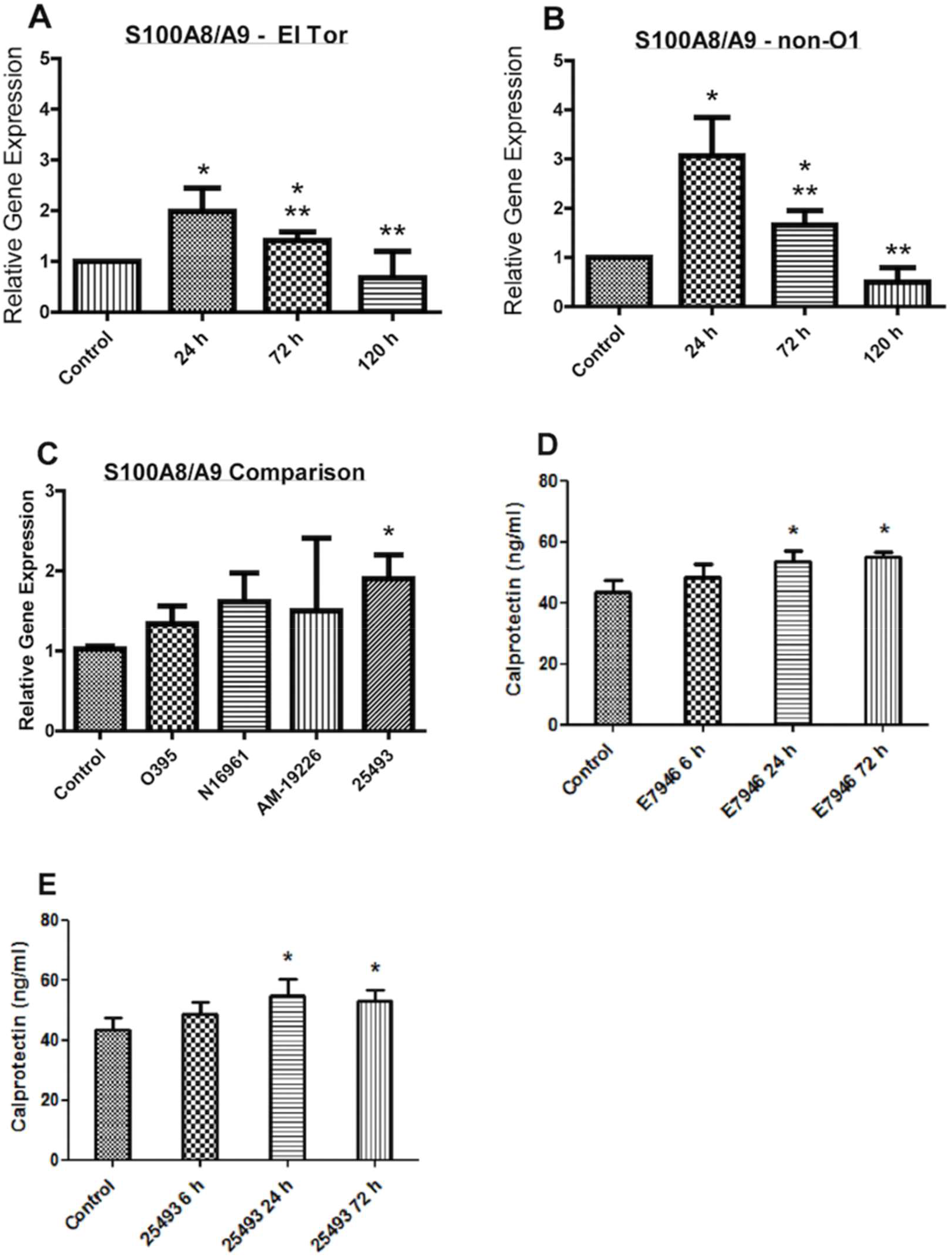
S100A8/A9 (calprotectin) levels increased during *V. cholerae* infection. WT zebrafish were infected with E7946 (El Tor) or 25493 (non-O1/O139) strains of *V. cholerae* at 2.5 × 10^7^ CFU/mL and then sacrificed at the indicated time points. mRNA levels were determined through qRT-PCR (3A-3C). Gene expression was normalized against β-actin and expressed as fold change. Protein levels were determined via ELISA (3D and 3E). Error bars indicate standard deviation. Data shown is from three experiments. *, P < 0.05 as compared to control, ** P < 0.05 as compared to 24 h infection.

Next, we determined the intestinal concentrations of calprotectin protein responses in the fish, as elevated fecal calprotectin levels are used as assays in a variety of disease states as being indicative of intestinal inflammation (18-20). For this experiment, we included a 6 h time point in addition to the significantly increased RNA time points of 24 h and 72 h. At the 6 h time point CP was not significantly increased, but at the 24 h and 72 h time points CP was significantly increased in fish infected with either of the strains, as was seen in RNA levels (Fig. 3D and 3E). Attempts to measure fecal calprotectin levels from these experiments proved unsuccessful (data not shown). This is likely due to the nature of the infection in the zebrafish, as the diarrheal fecal content is greatly diluted in the infection beaker water and could not be overcome by re-concentration.

### In-vitro effects of calprotectin protein on bacterial growth

Though the direct inhibitory effects of calprotectin have been shown for several other bacteria (17), no investigation has been done to date exploring its effects on *V. cholerae*. Therefore, we added increasing concentrations of purified calprotectin protein to cultures of *V. cholerae* strain 25493 in LB broth media, which proved to have dose-dependent growth inhibition, with complete inhibition of growth at 100 µg/mL (Fig. 4A).

**Figure 4.**
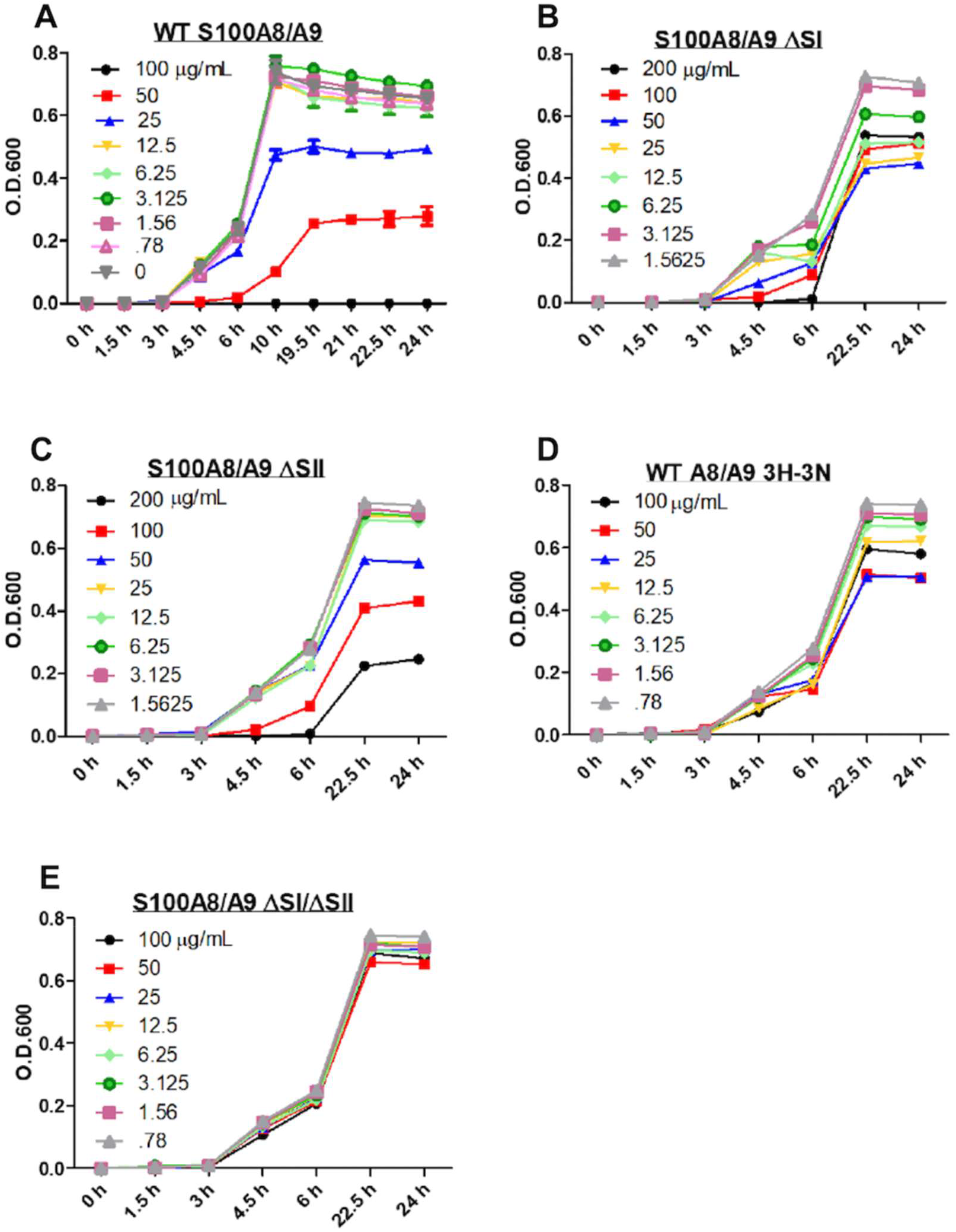
Zinc and Manganese sequestration inhibit *V. cholerae* growth. *V. cholerae* strain 25493 was grown in LB broth media with aeration at 37° C and measured at an O.D.600 at indicated time points. Increasing concentrations of WT (4A), ΔSI (4B), ΔSII (4C), WT A8/A9 3H-3N (4D), or ΔSI/SII (4E) calprotectin were added to LB broth media and incubated for 1 h prior to addition of bacteria. Data shown is from three independent experiments.

To further dissect how calprotectin exerts it anti-microbial activity against *V. cholerae*, CP mutants with altered binding site activity were used to explore the transition metal requirements for *V. cholerae* growth. The CP heterodimer of S100A8 and S100A9 have a pair of transition metal binding sites at the dimer interface (29). The unique site 1 (SI) is composed of six histidine (his) residues that have been shown to have high affinity for both Zn and Mn (17, 50). The canonical S100 protein site 2 (SII) is composed of 3-His, 1-Asparagine (asp) S100 protein that has high affinity for Zn but not Mn. CP variants used in this study include: (i) a knockout of binding by site II (ΔSII), leaving binding site I available to chelate Zn and Mn, (ii) a knockout of binding site I (ΔSI), leaving binding site II available to chelate Zn but not Mn; (iii) a selective Mn knockout (WT A8/A9 3H-3N) that substitutes two key S100A9 His residues for Asn in S2, leaving a tetravalent 4-His site with normal stoichiometry and high affinity for Zn; (iv) a site I and site II double knockout (ΔSI/ ΔSII) that lacks high affinity for transition metals, which serves as a negative control (17). Note that for accurate comparisons of growth inhibition, the concentrations of the ΔSI and ΔSII mutants were doubled relative to wild type CP because each dimer could only bind one ion instead of two.

The time course and concentration dependence of the inhibition of the growth of *V. cholerae* by CP was first investigated for wild type CP. Inhibition of growth was clearly evident at a CP concentration of 25 μg/mL and was complete at 100 μg/mL (Fig. 4A). No growth decreases were observed in the double binding site (Site I/II) knockout (Fig. 4E), consistent with growth inhibition arising from sequestration of transition metals. Assays with the selective Mn knockout mutant resulted in a decrease of bacterial growth by roughly one-third at the highest 100 μg/mL concentration (Fig. 4D), indicating an important role for manganese in *V. cholerae* growth. This was confirmed in the experiments with CP ΔSI, which also cannot bind Mn, as the level of growth inhibition was about the same at the highest 200 100 μg/mL concentration (Fig. 4B). Interestingly, the CP ΔSII inhibited bacterial growth by roughly two-thirds (Fig. 4C). This mutant retains the ability to bind Zn and Mn and only differed noticeably from WT CP only at the three highest concentrations. This difference could arise from an error in protein concentration or differences in solubility of the mutant versus wild type. Given that it can chelate both transition metals, we have been unable to conceive of other explanations for why this mutant is not ultimately as effective as the wild type protein. Regardless of this apparent anomaly, these results show that transition metals zinc and manganese both play important roles in the growth dynamics of *V. cholerae* and are essential for maximum growth potential.

### Growth recovery in the presence of calprotectin proteins

To confirm the antimicrobial activity of calprotectin was due to the chelation of these transition metal ions, we then grew bacterial cultures in the presence of CP and CP variants, along with increasing concentrations of Zn, Mn, and Zn + Mn in combination. Sub-inhibitory concentrations of wild-type CP at 50 µg/mL and 100 µg/mL of CP mutants was used. In the case of Zn addition, complete or nearly complete growth recovery could be seen for all three calprotectin proteins (Fig. 5A), as was the case for the Zn + Mn additions (Fig. 5C). However, the Mn only addition resulted in full growth recovery in only the wild-type CP and ΔSII, but not the ΔSI mutant, as was expected (Fig. 5B). Overall, these results confirm that the antimicrobial activity of calprotectin in limiting *V. cholerae* growth is through the sequestration and chelation of zinc and manganese.

**Figure 5.**
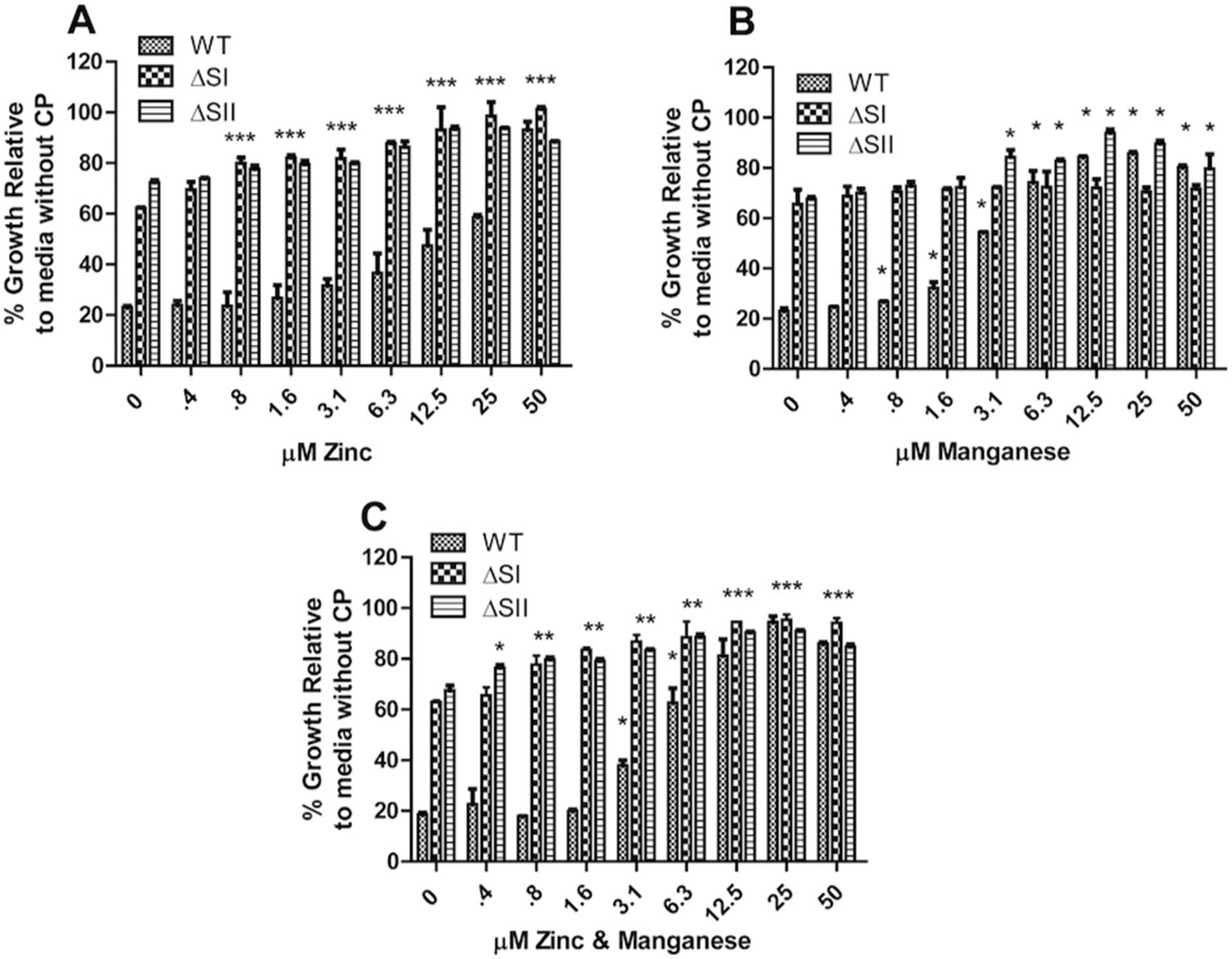
Zinc and Manganese are essential for maximum growth potential of *V. cholerae. V. cholerae* strain 25493 was grown in LB broth media with aeration at 37° C and measured at an O.D.600 at a 24 h time point. Sub-inhibitory concentrations of WT (50µg/mL), ΔSI (100µg/mL), or ΔSII (100µg/mL) calprotectin were added to LB broth media and incubated for 1 h prior to addition of bacteria. Zn, Mn, or Zn & Mn were added in increasing concentrations as indicated. Data shown is from three experiments and represented as % relative to bacterial growth in LB broth media alone. Error bars indicate standard deviation. *, P < 0.05 as compared to no metal addition.

### Is transition metal ion associated antimicrobial activity specific to calprotectin?

We then asked whether the antimicrobial activity of transition metal chelation was specific to calprotectin, or if this growth inhibition could be achieved via other metal limiting methods. The high affinity transition metal chemical chelator *N,N,N’,N’-*tetrakis (2-pyridinylmethyl)-1,2-ethanediamine (TPEN) was added directly to bacterial cultures at increasing concentrations and read at an O.D.600 at 24 h. This treatment resulted in complete inhibition of growth at high enough concentrations (Fig. 6A). As with CP, we next asked whether this TPEN growth inhibition could be overcome by adding transition metals Zn, Mn, and Zn + Mn in combination back into bacterial cultures. Using sub-inhibitory concentrations of TPEN (31 µM), nearly complete growth levels were recovered using all three combinations of metals (Fig. 6B). This confirms that limiting the availability of transition metals Zn and Mn has a direct growth inhibition effect on *V. cholerae*.

**Figure 6.**
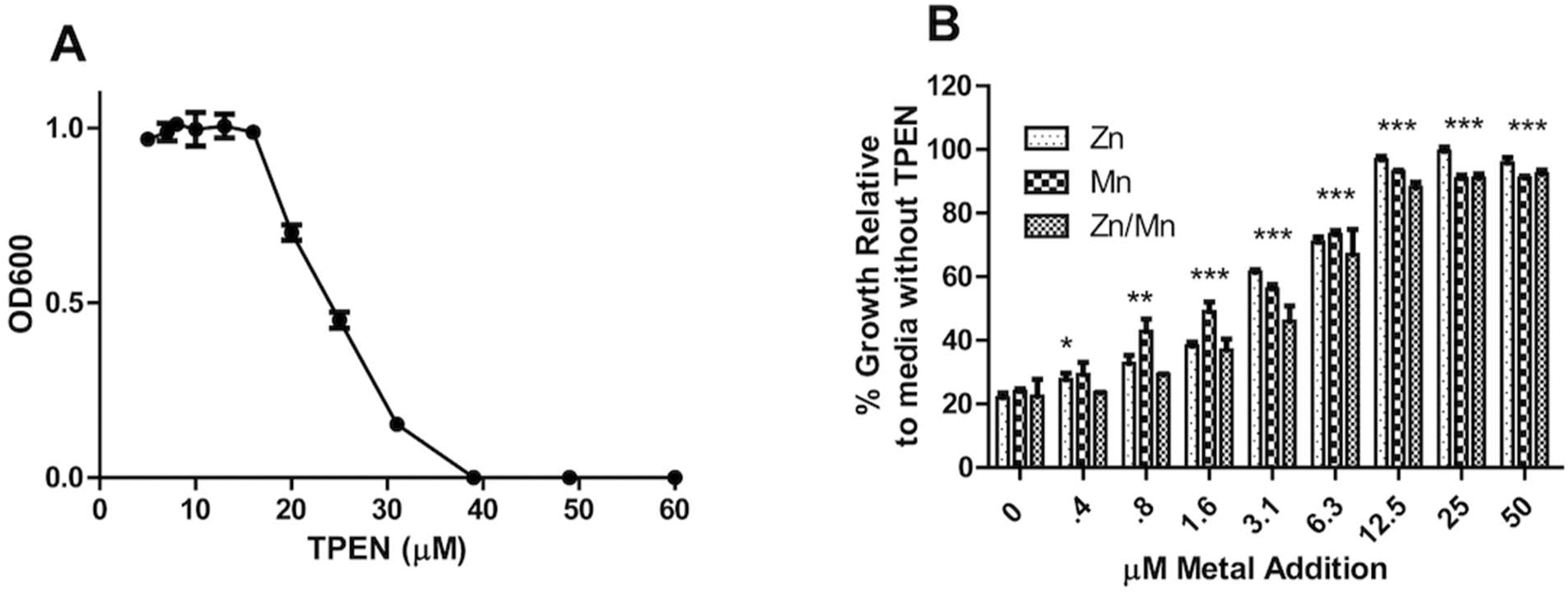
TPEN sequestration of Zinc and Manganese inhibits *V. cholerae* growth. TPEN was added to LB broth and incubated for 1 h prior to addition of bacteria. *V. cholerae* strain 25493 was grown in LB broth media with aeration at 37°C and measured at an O.D.600 at 24 h time points. Increasing concentrations of TPEN were added to culture (6A). A sub-inhibitory concentration of TPEN (31 µM) was added to LB broth media with increasing concentrations of Zn, Mn, or Zn & Mn. Data shown is from three experiments and represented as % relative to bacterial growth in LB broth media alone. Error bars indicate standard deviation. *, P < 0.05 as compared to no metal addition.

### Downstream effects of calprotectin in the zebrafish model

In addition to having antimicrobial effects, calprotectin is also well known to have immunomodulatory effects. Known as a damage-associated molecular patterns (DAMPs), CP along with other members of the S100 family can interact with pattern recognition receptors (PRRs) such as Toll-like receptors (TLRs) and RAGE (31). It can also initiate pro-inflammatory and anti-inflammatory responses, as well as act as a chemokine and cytokine. As an endogenous agonist of TLR4, CP binding initiates a cascade of signaling through the NF-κB pathway, leading to a proinflammatory response and neutrophil recruitment (32, 33). Direct lipopolysaccharide (LPS) binding has also been shown to be a potent activator of inflammation through TLR4 (35). Furthermore, calprotectin can act as a direct chemoattractant for neutrophils, leading to adhesion at the site of infection (34). Ultimately, calprotectin contributes to a positive feedback loop through pro-inflammatory signaling cascades that leads to active inflammatory responses. The zebrafish model was used to see if these components of inflammation are upregulated in response to *V. cholerae* infection. Both El Tor and environmental strains induced significantly increased levels of TLR4 and NF-κB in the fish at all three time points (Fig. 7A-D). Total leukocytes in the intestine of the zebrafish were also measured to gauge levels of leukocyte recruitment to the site of infection, with all three time points showing significant increases in response to either of *the V. cholerae* strains (Fig. 7E and 7F). Though not specific to CP alone, these results indicate that molecules known to be associated with CP are in fact increased in zebrafish, likely in part due to the immunomodulatory effects of calprotectin.

**Figure 7.**
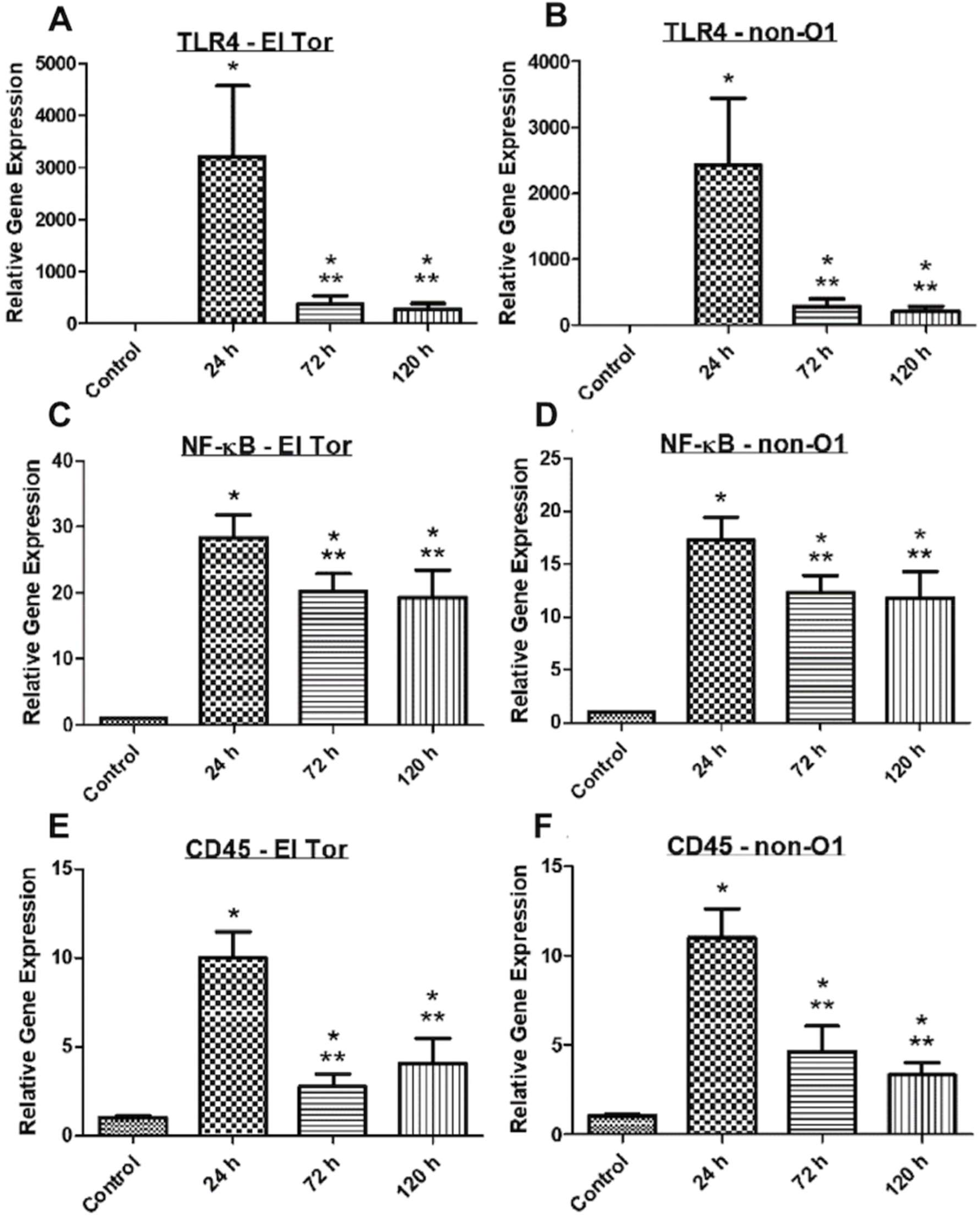
TLR4, NF-κB, and total leukocyte (CD45) levels increased during *V. cholerae* infection. WT zebrafish were infected with E7946 (El Tor) or 25493 (non-O1) strains of *V. cholerae* at 2.5 × 10^7^ CFU/mL and then sacrificed at the indicated time points. mRNA levels were determined through qRT-PCR. Gene expression was normalized against β-actin and expressed as fold change. Error bars indicate standard deviation. Data shown is from three experiments. *, P < 0.05 as compared to control, ** P < as compared to 24 h infection.

### IL-8 is essential in neutrophil recruitment

Finally, the importance of IL-8 in neutrophil recruitment and subsequent calprotectin release during *V. cholerae* infection in zebrafish was assessed. Previous studies have demonstrated the importance of IL-8 in recruiting neutrophils to the site of injury in the zebrafish (21). To achieve this, we used the selective CXCR2 (IL8R) chemokine receptor antagonist SB 225002 for drug inhibition at a concentration of 5 µM. We also utilized a splice blocking morpholino (MO) of cxcl8-l1 E1/I1 injected into single-cell stage embryos as an alternative method (21). In the drug Treated + Infection group, IL-8 and neutrophil RNA levels were significantly reduced as compared to the Untreated + Infection group, yet were still significantly increased as compared to the control Untreated – Infection group (Fig. 8D-8F). Calprotectin levels in the drug Treated + Infection group were significantly reduced as compared to Untreated + Infection group, while not having a statistically significant difference in levels as compared to the control Untreated – Infection group. Partial decreases in IL-8 and neutrophil RNA levels may be due to an inability of SB 225002 to completely outcompete IL-8 binding to CXCR2. Calprotectin RNA levels returning to basal control levels after treatment are likely a testament to the fact that a large amount of CP released in the gut is neutrophil derived. In cxcl8-l1 E1/I1 MO experiments, IL-8 splice blocking led to significantly decreased levels of IL-8, neutrophils, and calprotectin in the Treated + Infection group as compared to the Untreated + Infection group, while levels of neutrophils in the Treated + Infection group were also significantly decreased as compared to the control Untreated – Infection group (Fig. 8A-8C). These results highlight the importance of IL-8 in recruiting neutrophils to the site of infection, and the subsequent release of calprotectin.

**Figure 8.**
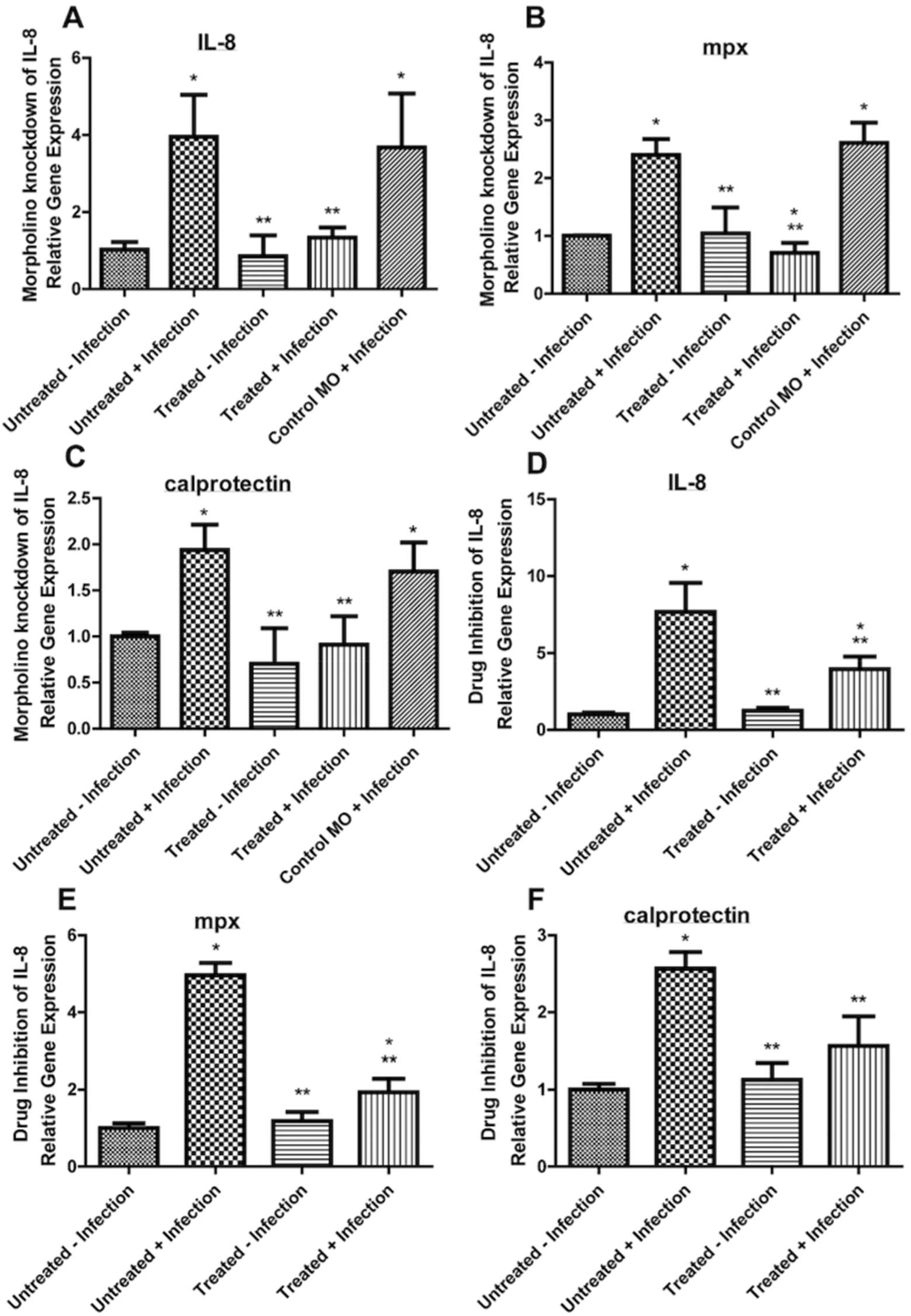
IL-8, neutrophils (mpx), and S100A8/A9 (calprotectin) levels decreased during *V. cholerae* infection in groups treated with morpholino (MO) (8A-8C) or drug (SB 225002) (8D-8F) (Treated + Infection). WT larval zebrafish were infected with 25493 (non-O1) strain of *V. cholerae* at 2.5 × 10^7^ CFU/mL and then sacrificed at 24 h time point. mRNA levels were determined through qRT-PCR. Gene expression was normalized against β-actin and expressed as fold change. Error bars indicate standard deviation. Data shown is from three experiments. *, P < 0.05 as compared to control Untreated - Infection, ** P < 0.05 as compared to Untreated + infection.

## Discussion

Here we describe the innate immune response of the zebrafish to *V. cholerae* infection, with particular interest in neutrophils and neutrophil associated molecules, as well as downstream markers of inflammation such as TLR4 and NF-κB. This work aids in further solidifying the already established utility of the zebrafish model in studying *V. cholerae* infection, as well as expanding its effectiveness into immune responses to *V. cholerae*, and the field of infection immunology as a whole. Additionally, new relationships between the nutritional immunity protein calprotectin and *V. cholerae* are established in this work, expanding its potential as a diagnostic marker or therapeutic target of interest.

Neutrophil responses in the gut have been reported to peak at 24 h and contribute to the intestinal inflammation that is seen in many GI diseases (28). In the current study, peak levels of neutrophil RNA at 24 hours in zebrafish were also observed (Fig. 1A and 1B), which coincides with previously seen peaks of *V. cholerae* levels in zebrafish intestines (23, 24, 37). Intestinal infiltration was also witnessed via fluorescent microscopy, where neutrophils could be seen in *V. cholerae* biofilms (Fig. 2B and 2C). At the site of infection, neutrophils are known to release extracellular traps (NETs), which are condensed chromatin that contains antimicrobial proteins to entrap and promote extracellular killing of pathogens. It has been reported that *V. cholerae* induces NET formation and release upon contact with neutrophils, and that *V. cholerae* utilizes two extracellular nucleases to degrade and evade these NETs (38, 39). Though the current study did not further investigate the relationship between zebrafish neutrophils and *V. cholerae*, future experiments similar to those seen here may provide further insight into this relationship.

One antimicrobial protein located on NETs is the nutritional immunity protein calprotectin. The antimicrobial activity of CP is well established, as is the mechanism through which CP inhibits microbial growth by transition metal chelation. Previously only Ellis et al. have shown an increase in S100A8 protein during infection in cholera patients (27); however, no other work has been done to explore this relationship. In the current study, RNA and protein levels of calprotectin were observed to be significantly increased at 24 h and 72 h in fish infected with either *V. cholerae* El Tor or the non-O1/O139 environmental strains and returned to baseline levels by 120 h (Fig. 3A and 3B), providing new evidence of this host response during infection. After showing that purified CP protein was able to completely inhibit *V. cholerae* growth (Fig. 4A), we then established that it does so in part through the chelation of the transition metals zinc and manganese (Fig. 5A-5C). Further evidence of this effect was also shown using the chemical metal chelator TPEN (Fig. 6A and 6B). It is well known that metals are tightly controlled and regulated by many biological systems, including *V. cholerae*, which uses metal transporters to tightly regulate intracellular levels (40). However, to our knowledge, this work shows for the first time the ability to inhibit *V. cholerae* growth through transition metal limitation. Further exploration into our understanding of how metal limitation inhibits growth may provide insight into exploiting this relationship as a potential target for therapeutics.

CP is also known to have downstream effects on the immune system. Using the zebrafish model, we showed that *V. cholerae* induces a significant CP response that leads to significant increases in the downstream receptor TLR4 (Fig. 7A and 7B), which then signals through the transcription factor NF-κB, a major regulator of both innate and adaptive immunity, and was also significantly increased in the fish model (Fig. 7C and 7D). This signaling pathway then leads to a positive feedback loop, amplifying the immune response. This in turn leads to more neutrophil and leukocyte recruitment, as does CP in its inherent chemokine activity, which can be seen in increased CD45 RNA levels in fish intestine (Fig. 7E and 7F).

The data in this study also show the importance of the cytokine IL-8 in the immune response to *V. cholerae* infection. By using morpholinos and drug inhibitors, IL-8 was found to be essential in neutrophil recruitment, and ultimately the release of calprotectin (Fig. 8A-8F). While the splice blocking IL-8 morpholino prevents the production of the cytokine IL-8, the drug SB 225002 works as a potent chemokine receptor antagonist that inhibits IL-8 from binding its receptor CXCR2 (IL-8R), despite increased levels of IL-8 still being produced in response to infection. Using the IL-8 morpholino, IL-8, neutrophils, and calprotectin all returned to baseline levels after MO treatment. Drug inhibition of CXCR2 did result in significantly decreased IL-8, neutrophils, and calprotectin as compared to the Untreated + Infection group, however they were still significantly increased as compared to the control Untreated – Infection group. Because IL-8 is still produced in this experiment, it may be outcompeting the drug binding to its receptor CXCR2. In future experiments, increasing amounts of SB 225002 drug treatment may ameliorate these issues. De Oliveira et al. previously described that IL-8 was upregulated in zebrafish in response to acute inflammatory stimuli and was ultimately essential for neutrophil recruitment and resolution of inflammation (21). Our data corroborate this, as IL-8 proved crucial in not just neutrophil recruitment, but also in calprotectin release, which comes primarily from neutrophils (13).

Use of the zebrafish model in studying *V. cholerae* infection is still relatively novel compared to other more commonly used mammalian animal models, however its advantages are numerous. Among them are the availability of transgenic animals, transparent embryos, and relatively lost cost, as well as several others (3). The work done here continues to strengthen the evidence of using this model in studying *V. cholerae* infection. In addition, this study expands the zebrafish model to the field of *V. cholerae* immunology, where a number of questions remain unanswered, such as how different strains of cholera cause differing levels of inflammation, how to create more efficacious vaccines, and how the gut microbiome affects pathogenesis (41).

Prior to this study, no published work had assessed the zebrafish calprotectin response. One reason for this may be due to a general lack of immunology related tools in the zebrafish field, as compared to other animal models. The data in this study provide evidence that the zebrafish is capable of mounting a calprotectin response, similar to those seen in humans and other mammals. For the first time a link between nutritional immunity and *V. cholerae* has also been established. The data presented here solidifying calprotectin responses to cholera calls for more investigation into this relationship, as well as other antimicrobial nutritional immunity proteins such as S100A12 (calgranulin C), which has been shown to be upregulated in response to other infectious diseases (42-44). One area of interest is the possibility of calprotectin as a diagnostic marker. Though cholera can be diagnosed through rapid dipstick methods or simply diagnosed through symptoms in endemic areas, CP may provide another alternative for diagnosis and serve as an indicator of levels of inflammation in the patient. Due to its stability (stable in feces up to 1 year when frozen), CP may be useful in epidemiological studies where samples must be revisited at a later date. Drawbacks to using this protein for diagnosis remain in that intestinal calprotectin release may not be specific for differentiating cholera from other bacterial infections or other inflammatory GI diseases, which can have increases as large as ten-fold and exceed 1 mg/ml (45, 48). In addition, the relationship between certain bacteria and CP needs further exploration, as the presence of CP can enhance some bacterial growth (46, 47).

In summary, the data presented here provide new approaches for the *V. cholerae* field using calprotectin as a potential diagnostic marker or therapeutic. This work has also established the ability to inhibit *V. cholerae* growth through metal limitation, giving further evidence that transition metals are essential for *V. cholerae* growth. The ability to alter or limit transition metal distribution and homeostasis in infected tissues may be an effective method in antibacterial development. Complete growth inhibition by CP would likely take excess metal binding capacity, due to competition with high affinity bacterial metal uptake systems (30). Further illumination of the pathogen factors affected, their intrinsic nutrient requirements, and the complex interplay between host and pathogen factors have the potential to open new avenues for the development of anti-*V. cholerae* therapeutics.

## Ackowledgements

Many thanks to past and present members of the Withey lab for helpful discussions. This work was supported by Public Health Service grant R01AI127390 (to JHW).

